# MS2AI: Automated repurposing of public peptide LC-MS data for machine learning applications

**DOI:** 10.1101/2021.01.27.428375

**Authors:** Tobias Greisager Rehfeldt, Konrad Krawczyk, Mathias Bøgebjerg, Veit Schwämmle, Richard Röttger

## Abstract

**Motivation:** Liquid-chromatography mass-spectrometry (LC-MS) is the established standard for analyzing the proteome in biological samples by identification and quantification of thousands of proteins. Machine learning (ML) promises to considerably improve the analysis of the resulting data, however, there is yet to be any tool that mediates the path from raw data to modern ML applications. More specifically, ML applications are currently hampered by three major limitations: (1) absence of balanced training data with large sample size; (2) unclear definition of sufficiently information-rich data representations for e.g. peptide identification; (3) lack of benchmarking of ML methods on specific LC-MS problems.

**Results:** We created the MS2AI pipeline that automates the process of gathering vast quantities of mass spectrometry (MS) data for large scale ML applications. The software retrieves raw data from either in-house sources or from the proteomics identifications database, PRIDE. Subsequently, the raw data is stored in a standardized format amenable for ML encompassing MS1/MS2 spectra and peptide identifications. This tool bridges the gap between MS and AI, and to this effect we also present an ML application in the form of a convolutional neural network for the identification of oxidized peptides.

**Availability:** An open source implementation of the software can be found freely available for non-commercial use at https://gitlab.com/roettgerlab/ms2ai.

**Supplementary information:** Supplementary data are available at *Bioinformatics* online.

## 1 Introduction

LC-MS is the leading technique to measure the cellular proteome. LC-MS is routinely performed as tandem-MS, as this procedure allows in-depth identification of the proteome, and delivers information such as sequence and post-translational modifications based on a database search of the secondary mass spectra (MS2)^1^. When analyzing tandem-MS data, the full information of primary peptide mass spectra (MS1) and MS2 is mostly simplified, and no current pipeline converts the full wealth of data into a format amenable for ML. In addition, the plethora of publicly available MS data is a mostly untapped resource for advanced ML techniques. It is thus desirable to establish a connection between available LC-MS data and ML applications in order to unravel any latent but biologically or technically relevant information. Advanced ML methods like deep learning models have shown great potential to predict peptide features of different charge states, as well as estimate their intensity^2^, tumor classification using imaging mass spectrometry^3^ and peptide MS2 spectra prediction^4^.

To address the issue of data connectivity, we created MS2AI, a pipeline ensuring that advanced ML techniques are applicable to large scale MS data. This is done by standardizing heterogeneous data resources for reliable repurposing, and further enrichment with experimental and peptide-centric metadata. MS2AI solves the fundamental challenges of ML in MS suffering from the lack of convenient acquisition of large-scale training and test data. MS2AI automatically extracts compatible entries (analyzed with MaxQuant) from the largest public repository of LC-MS data, PRIDE^5^, and stores these in an homogenous and ML ready standardized format.

## 2 Methods

### 2.1 Automated data retrieval

MS2AI fetches metadata from all ∼12.000 PRIDE projects using the PRIDE API and stores it in a MongoDB collection. These projects can be filtered according to specific metadata entries (see supplementary paper table 1). For convenience, MS2AI contains a complete database collection of PRIDE project metadata (as of January 202) that is easily updatable to include newer projects.

Currently, MS2AI can only process projects that come with peptide identifications from the MaxQuant suite, which is the predominantly used software for bottom-up proteomics. This information yields details about peptide identity and quantification, such as the peptide location, sequence, intensity and modifications. By restricting the database to MaxQuant identifications, MS2AI ensures a more homogenous identification and extraction of peptides across all projects and their raw data files.

Based on the PRIDE metadata file, which is automatically filtered with regards to MaxQuant analysis and user-specified requirements, MS2AI retrieves the MaxQuant output files, along with the corresponding raw files from the PRIDE database. These raw files are in a proprietary file format used by the majority of MS instruments in PRIDE, and contain information from the data acquisition process, most notably the objects in the m/z–retention time–intensity space^6^.

The following data transformations are executed after raw file download; (1) conversion of raw files into the community standard mzML data format using the ThermoRawFileParser^7^, and (2) further extraction from the mzML file, including all MS2 spectra, and values in the m/z–retention time–intensity space from the MS1 spectra. The output of the processed raw files are stored in the local data directory, while the raw files themselves are discarded (see supplementary data for file storage structure).

Along with the automatic PRIDE data retrieval, MS2AI also allows for the extraction of local in-house data. This requires the MaxQuant and corresponding raw file(s) to be available to the software. The key steps of the extraction are identical to the steps taken during the PRIDE method. The user has the possibility to provide additional metadata to their local data comparable to the metadata (instrument, modifications, etc.) available on PRIDE to seamlessly integrate local and PRIDE-received data.

### 2.2 Peptide representation

From the extracted information described in section 2.1, we are able to create a run representation (RR) of the entire LC-MS run (see figure 1). However, due to the sheer amount of m/z and retention time entries in the LC-MS spectra, data reduction is paramount, as LC-MS base resolution would make ML methods insurmountably computationally challenging (see supplementary data Figure 6). The user can freely define the range of m/z and retention time to be summarized into a single data point of the RR; the smaller the range in both m/z and retention time, the less data loss will occur, at the expense of increased data size. This increase in size has an effect on both storage space and the runtime of any future ML applications (see supplementary data section 6). To account for the inevitable data-loss on the RR, we have constructed a 4-channel data representation of the MS1 spectra consisting of: (1) mean value of all summarized data-points, (2) minimum value of all summarized data-points, (3) maximum value of all summarized data-points, (4) absolute number of summarized data-points. To avoid unwanted bias in the intensities, all intensities in a single raw file are normalized based on the highest value in the individual raw file. The peptide representation (PR) consisting of the peptide and the neighbourhood, which size is configurable, are drawn directly from RR, along with the MS2 spectra for the corresponding peptide.

**Figure 1.**
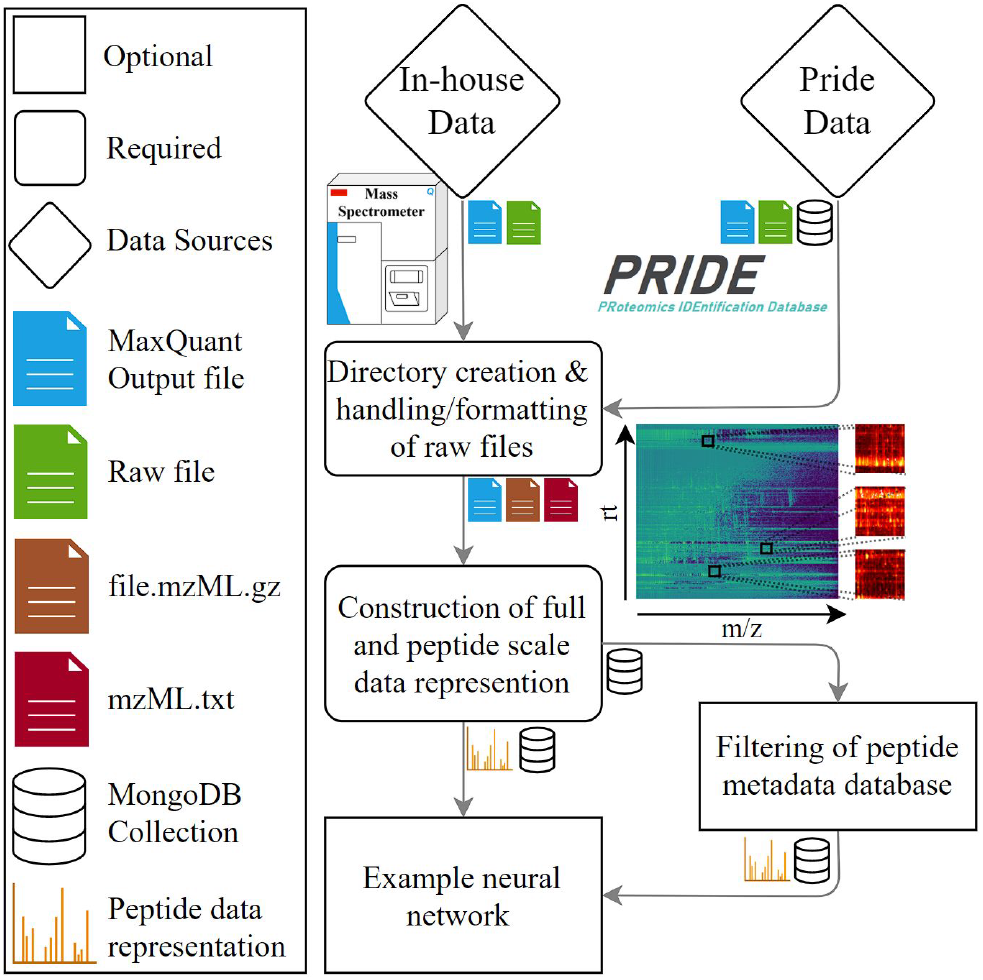
Pipeline structure and workflow, with legends explaining what files are transferred to future parts of the workflow, and a large scale image of the data representation of MS1 data.

The MS2 information is either extracted exactly as represented in the mzML file (with differing length of m/z-intensity space), or in a binned fashion, to ensure equal length and homogeneity between spectra. The m/z bin-size of MS2 can be calculated in two methods; fixed length (e.g 500 bin in 0-2500 m/z) or in a variable length (e.g. 500 bins in 0-precursor m/z). Furthermore, intensity information of bins can be kept as the mean of normalized values, or dichotomized indicating whether or not a peak exists in the binned area. All of these options give users full flexibility to adjust the PR to their specifications.

Each PR is accompanied by all relevant information from the MaxQuant output file, which is appended to a MongoDB collection for machine learning purposes. This collection can then be filtered and sorted to help transitioning into any ML application, by changing or removing entries based on needs.

### 2.3 Machine learning applications

The MS2AI PR is well suited for a multitude of ML applications, as it delivers homogenous data representations of heterogeneous MS experiments. The generation of PRs, along with the metadata accompanying each of the peptide data, trivialize the task of getting from unsuitable raw data to ML ready peptide information files.

MS2AI also includes a functional convolutional neural network that uses a tailored data generator for TensorFlow^8^, allowing easy integration of the 4-channel images along with m/z and rt information from MS1 and the MS2 spectra. In order to demonstrate the utility of MS2AI, we trained and tested a simple neural network (see supplementary information Figure 6) on ∼200.000 PRs from 307 different projects (filtered from ∼69.000.000 total PRs with 98th percentile score filter) separated into peptides with an oxidation on methionine, and peptides without an oxidation on methionine. We then trained and tested whether the network could, using MS1 and MS2 information, distinguish the two classes. Doing this we obtained 95% training accuracy and 93% validation accuracy along with 85% test accuracy on a different set of PRIDE projects consisting of 25.000 PRs; entire projects from which none of the data were not used for training or validation. This separation of training and test data causes highest possible heterogeneity between data points and robustness in the neural network capabilities (see supplementary information section 5).

## 3 Conclusion

MS2AI is the first automated pipeline for strategic and large scale extraction and processing of LC-MS experiments that allows hassle-free and powerful ML applications in the realm of computational proteomics. By combining the measurements in the multidimensional area around a peak on the m/z–retention time–intensity space with the fragmentation MS2 spectra. MS2AI also offers unique concise peptide data representation that contain the most vital MS1 and MS2 information to describe a given peptide; m/z values, retention times and measured intensities. This PR is highly customizable and thus enables researchers to summarize peptide information in accordance to specific needs, along with a comprehensive database structure of known information on the peptides gathered from MaxQuant.

In general MS2AI will allow powerful applications of ML techniques performed in the field of MS, opening the door for more in-depth analysis of the proteome, as demonstrated by our example network identifying peptide modifications.

## Acknowledgements

The authors want to thank Alex Gherega for his valuable feedback and help to structure the software and streamline the development process according to industry standards.

