## supplementary data for "MS2AI: Automated repurposing of public peptide LC-MS data for machine learning applications"

*Data and text mining*

#### *Supplementary Paper*

Tobias Greisager Rehfeldt[1], Konrad Krawczyk[1], Mathias Bøgebjerg[1], Veit Schwämmle[2*], Richard Röttger[1]

[1]Department of Mathematics and Computer Science, University of Southern Denmark, [2]Department of Biochemistry and Molecular Biology, University of Southern Denmark

**Contact:**

*Table of Contents*

**1. Pipeline structure** **2**

**2. Data sources** **4**

a. PRIDE 4

b. Local 5

**3. Data representations** **6**

a. Run representations 6

b. Peptide scale data representation 7

c. Metadata Database 9

**4. Data filtering** **9**

**5. Use case neural network** **10**

a. Data 10

b. Architecture 11

c. Metrics, cost functions and optimizers 12

d. Results 12

**6. Runtime evaluations** **12**

**7. Tutorial** **13**

a. Setup 13

b. Data extraction 13

c. Filtering 14

d. Machine learning application 14

### 1. Pipeline structure

The generation of data suitable for machine-learning (ML) purposes requires multiple steps to retrieve, extract, organize and convert the data. Figure 1 shows the different parts of the pipeline.

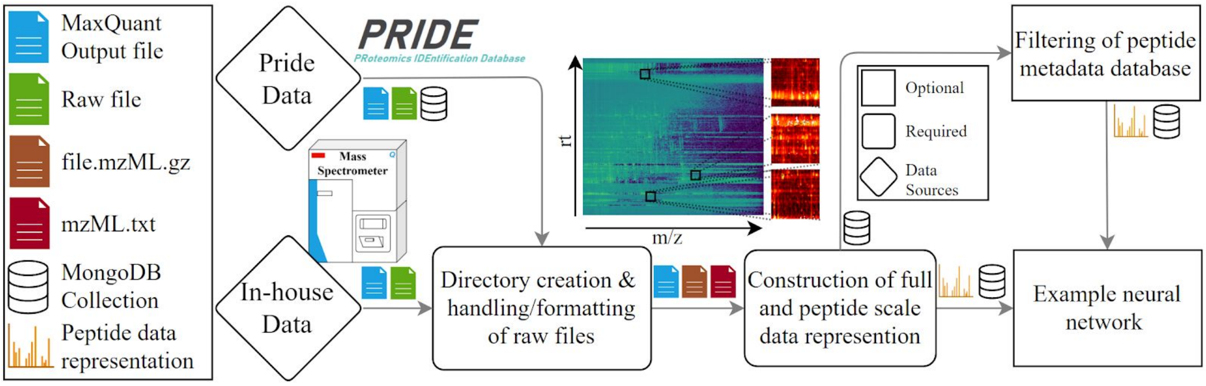


Figure 1. The structure of the MS2AI software pipeline

***Metadata and raw file collection***

The first part of the pipeline is concerned with the proteomics identification database (*PRIDE*) metadata and raw file collection. The collector gathers metadata of all projects in the PRIDE repository using the PRIDE API. The collector does not only retrieve all available information from the API interface, but also checks which files are available in the projects that mention MaxQuant as software used for quantitative data analysis.
The reason for checking the available files is that MS2AI relies on MaxQuant files for identifying the peptide locations (see section 3b). Projects that mention MaxQuant and have any of the suitable output files (allPeptides.txt and evidence.txt) will be amenable for MS2AI. Additional filtering options to the API metadata can be added, but are required to be directly included into the python script by the user itself. After automatically retrieving suitable PRIDE projects that fulfill the filter criteria, the next step is downloading and matching MaxQuant files with the corresponding Thermo raw files on the PRIDE database.

The current PRIDE metadata collection (as of January 2021) metadata file is available here. The MS2AI api also comes with an inbuilt method to import the current PRIDE database, and is performed by the code below in a terminal from the ms2ai directory. For further information on the PRIDE metadata web-scraper see *documentation.pdf*.

```
Python api.py --db
```

A full introduction to the MS2AI database structure layout, the parameter settings and the installation instructions are available in the README file on Gitlab (https://gitlab.com/roettgerlab/ms2ai).

***Data organization and folder structure***

The second step in the pipeline is to format and handle the files that have either been acquired from the PRIDE metadata, or provided from in-house data. This consists of setting up a local directory structure, and converting the raw files into a non-proprietary community-based mzML data format. The converted files are stored in the folder structure depicted in Figure 2.

Each project resides in its own folder, where projects refer to either the projects from PRIDE (defined by their ProteomeXchange identifier PXD) or any local in-house folder used to extract data. A project will have sub-directories for each raw file within the given project, and each raw file directory will contain specific files necessary to create the peptide data representations (PR), these are: MaxQuant file, the formatted mzML.gz file, the data file with extracted data from the mzML.gz file, and the LC-MS run data representation (RR) files. For details about the data representation, see Section 3.

```
/path/to/data
├ images
│   └database-name
│       └project-name
│           └raw files
│               └peptide-representation-ID.txt
├ metadata
│   ├best-model.h5
│   ├network-model.png
│   └historic-plot.png
├ PXD projects
│   └raw files
│       ├ file.raw
│       ├ file.mzML.gz
│       ├ mzML.txt
│       ├ MaxQuant-file.txt
└       └ XxY.txt
```
Figure 2. Folder structure created by the MS2AI software at the path specified in the configuration file.

The '*images*' directory contains all PRs, where each PR is saved as a unique ID based on the ms/ms scan number. The images are saved in sub-directories of 'database name', 'project name' and 'raw file name', as this structure is chosen to allow multiple databases of PRs to co-exist. The MongoDB (MS2AI database structure) collection contains the metadata attached to each peptide, and the database-name directory contains the peptide data itself. This allows for multiple PRs of the same peptide to be created, for example to test different PR sizes or binning methods.

The '*metadata*' directory keeps all the information about neural networks composition and results. This directory will keep historic performance plots of each neural network, as well as the best model gathered during training and an image representation of the neural network itself. This is the folder in which the pre-trained networks reside, both for inserting pre-trained networks or collecting user trained networks. The folder is created in the late stages of the pipeline when working with neural networks. Currently, only a deep learning model is provided with the MS2AI software, so other ML techniques have to be user generated.

***Raw data processing***

The next step in the pipeline is the raw data processing.. Each raw file is converted into a zipped mzML file (*file.mzML.gz*). This is done using either a Conda or a dockerized version of ThermoRawFileParser[1] (*TRFP*). Along with formatting the file, the user also has the ability to apply peak picking performed by the native Thermo packages during formatting, which helps sort out noise from signal. MS2AI then extracts all MS1 and MS2 spectra information into a concise and easily readable format used for RR and PR creation (*mzML.txt*) for subsequent data generation tasks.

***Data representations***

The next part consists of creating the data representations, both of the LC-MS RR and the individual PRs (section 3). The RR is saved in the raw file specific folders, inside the project folders (*res(m/z)-x-res(rt).txt*), and the PR are saved in the images directory for usage in ML applications (*peptide-representation-ID.txt*). This allows fast creation of new data sets for subsequent ML approaches.

***Filtering and preparation to machine learning***

Before the inbuilt example neural network can be utilized, the peptide metadata has to be filtered. The filtering process accepts arguments (for instance PTMs or instrument names) given in either the MaxQuant files[2] or the PRIDE API[3], and classifies them into a new collection based on these criteria (see section 4. for more information of classes and arguments). Additionally, the data can be filtered according to their identification score assigned by MaxQuant (set in the configuration file). Filtering for high-scoring peptides will ensure high confidence identifications, but it will also come at a cost of reducing the amount of data available after filtering.

The output of this filtering process is stored as a new MongoDB collection of peptide metadata within the given database. This filtered collection contains only the bare requirements for subsequent machine learning to reduce storage demands, these are information such as classification name, class, id and data size.

***Machine learning***

This pipeline structure allows easy transition from raw LC-MS data into a concise and powerful data format. Hence, MS2AI mediates the connection between mass spectrometry and advanced machine learning such as artificial intelligence applications (*MS & AI*). We simplified the process of training new neural networks, or to utilize pre-trained neural networks for usage on local data, by integrating a pre-built neural network in the library Keras and allowing easy integration between the neural network and the data created in the MS2AI software. The example neural network inbuilt into MS2AI is explained in section 6 on network architecture. It is important to note that the data prepared by MS2AI is also amenable to any other machine learning algorithms.

### 2. Data sources

MS2AI is a tool used for extracting data from resources containing raw LC-MS data files analyzed by the MaxQuant Suite. Peptide identifications and quantifications from other tools can be added by either converting them to the MaxQuant text file formats or adaptation of the code. Currently there exists a multitude of different online data repositories. For this reason, we have implemented two/three methods to feed the software with MS data; **a**. data is retrieved from the PRIDE database (suitable for large scale data generation) and **b**. integration of local resources like in-house data. We expect to extend these inputs to allow for a wider range of public repositories.

#### a. PRIDE

MS2AI gathers all the PRIDE projects along with the metadata available through the PRIDE API. This information is kept in a MongoDB collection called ‘pride’ within the ‘ms2ai’ database. This collection allows for easy filtering on any of the keys available in the project API json entry. The full list of keys and their descriptions available for filtering are given in Table 1.
The ‘pride’ database contains three additional entries: filetypes (.zip, .raw, etc.), a boolean value describing whether or not the project has been analyzed using MaxQuant, and a list of the available MaxQuant output files (given that the boolean value was true). These three keys are considered in the ‘base-filter’ when running the inbuilt PRIDE data retrieval method, which checks that they are compatible with the software. The rest of the keys in the list are usable as user-customizable filters adaptable to specific needs, this could for example be restrictions to certain post-translational modifications, certain features of the experimental protocol, or a specific instrumental platform experiment. User-specific filters can be realized by directly implementing them into the python scripts by modifying the pride_data function and the pride_filter variable in the extractor/run.py script.

| Name | Data Type | Description |
|---|---|---|
| doi | string, optional | the Digital Object Identifier (DOI) for the project (if available), |
| quantificationMethods | array[string], optional | the quantification method(s) used with the dataset (if any) |
| submissionDate | string, optional | the date the project has been submitted, |
| submitter | ContactDetail, optional | details of the submitter of the dataset, |
| keywords | string, optional | relevant keywords associated with the project, |
| reanalysis | string, optional | annotation to indicate that the dataset is a re-analysis based on other public data, |
| experimentTypes | array[string], optional | the type(s) of experiment performed, |
| references | array[Reference], optional | publications/references associated with the project, |
| labHeads | array[ContactDetail], optional | the Lab-Head or PI associated to the dataset, |
| sampleProcessingProtocol | string, optional | project meta-data: information about the sample processing, |
| dataProcessingProtocol | string, optional | project meta-data: information about the data processing, |
| otherOmicsLink | string, optional | links to other datasets related to this project (if available) |
| numProteins | integer, optional | project statistics: number of reported proteins, |
| numUniquePeptides | integer, optional | project statistics: number of unique peptides, |
| numPeptides | integer, optional | project statistics: number of reported peptides, |
| numIdentifiedSpectra | integer, optional | project statistics: number of identified spectra, |
| numSpectra | integer, optional | project statistics: number of spectra, |
| title | string, optional | the title given to the project, |
| accession | string, optional | the project's accession number, |
| publicationDate | string, optional | the date the project has been made public, |
| tissues | array[string], optional | the tissue annotation for the project, |
| ptmNames | array[string], optional | the Post Translational Modifications (PTM) annotated for the project, |
| projectDescription | string, optional | the description provided for the project, |
| numAssays | integer, optional | the number of assays associated with this project, |
| submissionType | string, optional | the type of submission (complete or partial) |
| projectTags | array[string], optional | specific tags added to the project for classification, |
| species | array[string], optional | the species annotation for the project, |
| instrumentNames | array[string], optional | the instrument annotation for the project |

Table 1. Table of PRIDE API[4] key names, along with the data type and the description of the key, as stated in the API.

MS2AI will only download data from projects that satisfy the applied filters. The base filter requires that projects to mention 'MaxQuant' (a proxy variable, as there is no api key for quantification software) and to have the specified (in the configuration file) MaxQuant output file in the maxquant file list (allPeptides.txt[5] or evidence.txt[6]). When using a modified version of the filter, it is important to be case and space sensitive, as MS2AI needs an exact string matching in order to properly match with the entries in the PRIDE database.

Please note that iterating through all suitable data from PRIDE requires ~80 TB of disk space and, depending on the connection and compute facilities, might require several weeks or months to complete. For this reason effective filtering prior to data collection is paramount.

In addition to running all PRIDE projects in the database, there are also options to extract data from single projects, or a user-defined list of projects. To do this, follow the instructions in the *documentation.pdf* in the MS2AI directory.

#### b. Local

The other method of feeding data into, is via a local data source. In order to gather sufficient data to train a neural network, large-scale in-house data is required, we recommend to use at least one hundred raw files. When the user wants to extract data from local raw files, is it required that the raw and MaxQuant (or a zip file containing the MaxQuant) files are in the same directory, and that this path is given to the MS2AI software (see *documentation.pdf*). There can be multiple raw and MaxQuant files in the same directory, in which case the software will take advantage of the multi-threading capabilities of the local machine (number of processes is user-defined in the configuration file). The files will then be moved into the directory specified in the 'path' variable in the configuration file, in the same fashion as PRIDE PXD projects, where the project-name becomes the name of the folder the data is located in, see Figure 2.

### 3. Data representations

MS data comes from high-throughput experiments and is rich in information. Such data that can be very difficult to handle, interpret and re-formulate into ML amendable formats. MS2AI uses its own novel data representation, by creating representations of both the entire LC-MS run (RR) and the individual peptides (PR). MS2AI does not automatically locate peptides in the MS1 nor the MS2 spectra, but instead utilizes the MaxQuant identifications  to match peptides found based on the MS2 spectra with the locations in the MS1 spectra. More specifically, MS2AI utilizes either the allPeptides.txt or the evidence.txt file from MaxQuant. It should be noted that based on which type is chosen, different projects will be available from PRIDE and that the peptide metadata will contain differing keys. It is possible to utilize both MaxQuant output types, although this requires special care for aligning the different metadata information.

One of the main features of MS2AI is the highly customizability of the data representations (described in subsections a. and b.). For this reason we've implemented a method for re-creating all previously processed raw files, to efficiently re-create PRs after changes have been made to the extraction parameters. Both the RR and the PR come with native machine resolutions in m/z and retention time for MS1 that would far exceed current computations, and for this reason MS2AI utilizes a 4-channel representation of this data, one channel for each: (1) mean value of intensities, (2) minimum value of intensities, (3) maximum value of intensities, and (4) number of collapsed values. In (1)-(3) the intensity values are normalized between zero and one, and in (4) the log values of the counts are used. The MS2 spectra are kept in the PRs, in either full form, or in a binned version (explained in section b).

All user-defined settings for creating these data representations can be found in the configuration file (*config.json*) under the 'extractor' key. All of the parameters for all parts of the software are explained in the README file under the section '*Configuration file'*

#### a. Run representations

Saving MS1 spectra at the machine resolution that MS instruments outputs would lead to extensive storage space requirements and slow computations for both creation and ML applications. For this reason one of the key aspects of extracting peptide data, is to find an efficient method of representing the MS1 spectras in a concise format, allowing it to be saved with lower storage facilities and faster computational speeds.

MS2AI uses an innovative approach to create heatmap-like files with retention time on the y-axis, m/z on the x-axis, and signal intensity as the corresponding values between the axis. Examples are shown in Figure 3, where an entire LC-MS run is divided into multiple 2-dimensional representations (combined into one higher dimensional representation).

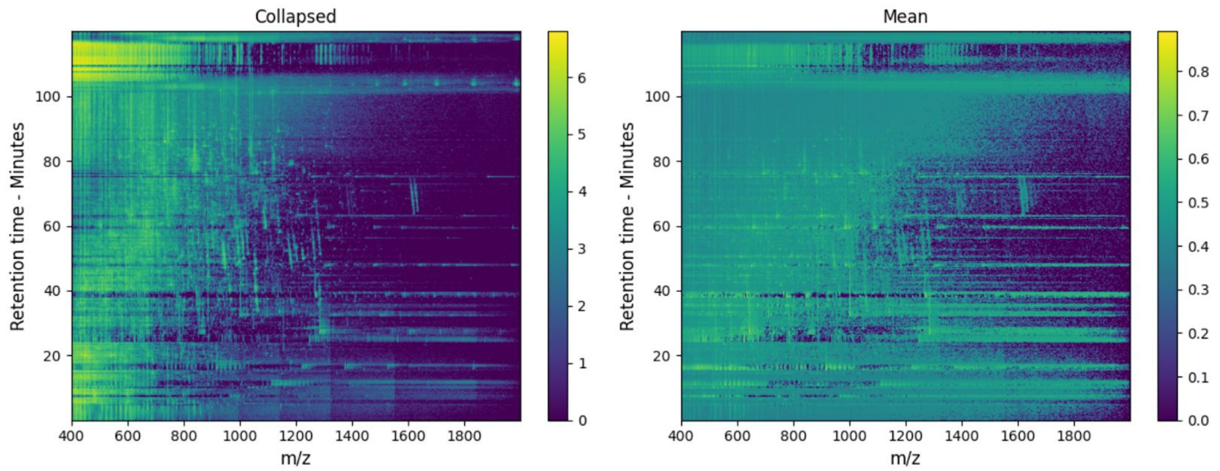


Figure 3. Image representation of two of the 4 channels available in the full scale data representations; mean intensity and amount of collapsed values.

As mentioned earlier, the machine resolution of the entire LC-MS run is far too large to handle during ML training and data creation, and furthermore the peak distribution is far too sparse to really derive any meaningful information of the surroundings. For these reasons we need to simplify the high resolution in the m/z and retention time space into single data-points while still maintaining as much information as possible. This was achieved by MS2AI's 4-channel version of the MS1 data.

This is done by collapsing a certain range of both m/z and retention time points into a singular data point (or pixel in the image). How many data points are collapsed into a single point is denoted by the bin sizes; A smaller bin size results in a larger and more detailed image, but also slows down the computations (see section 6 on runtime calculation). These bin sizes are configurable by the user (*config.json*), with defaults set in accordance with our test case (2 m/z pr bin and 0.2 rt pr bin). The bin size should be configured depending on what the study tries to utilize and predict based on the PRs, and the ML technique used.

#### b. Peptide scale data representation

Along with the RR, MS2AI creates concise and highly customizable PR. Here, the way the RR is constructed has a direct effect on the PR, as the PRs are directly extracted from the RR. Consequently, the size of the bins in the RR defines the size of the bins in the PR. Creating PRs directly from the RR is a computational compromise between speed and data precision. If every PR was calculated based on the mzML file, there would be higher levels of detail and customizability, but it also comes at a much higher computational cost, as identical areas would to be calculated multiple times for separate peptides.

Along with the compressed versions of MS1 data, the PRs also contain details of associated MS2 spectra. These can either be represented as full spectrum, or in a binned fashion to create a summarized and more general version.

The idea behind these PRs is that there will be a homogeneity in not just the analyzed peak, but also in the neighbouring peaks across separate RR for similar peptides, meaning that any given peptide has a generalizable neighbourhood of similar peaks. The important parameters for extracting PRs are the extracted neighbourhood size around the peptides (both m/z and rt) for the MS1 spectra, and the specific method used to simplify and balance the MS2 spectra. These parameters are set in the configuration file (*config.json*).

*MS1 level configuration.*

The size of the area surrounding the peptide peak in the RR is called the neighbourhood and can be defined in the *config.json* file. Large areas with high resolution result in slower computations - both in creation and ML applications (see section 6 on runtime calculations). An example of the PR is seen in Figure 4.

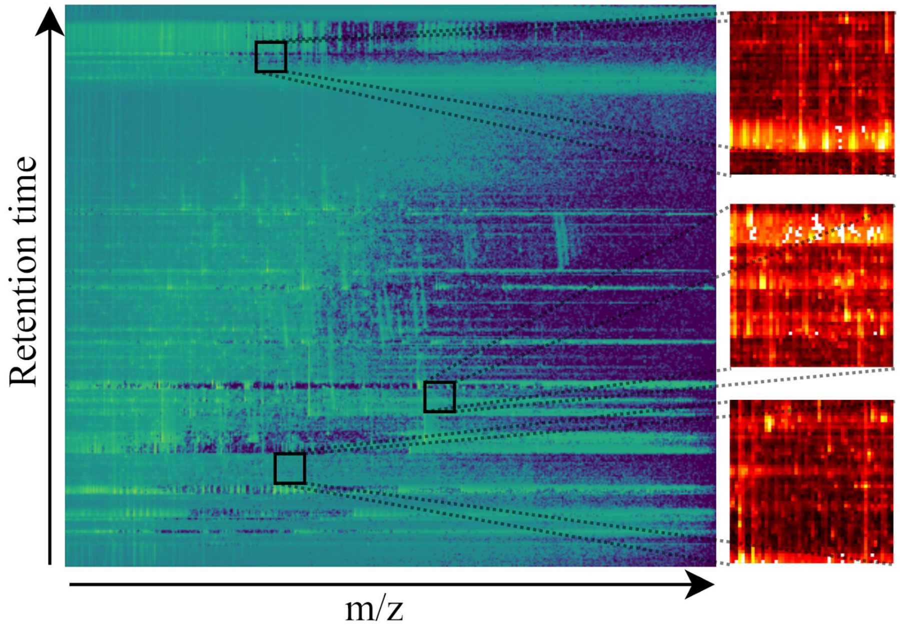


Figure 4. RR and PR from one of the 4 channels (mean). Two peptides are found in the full scale data and expanded into PR.

When extracting the peptide data from the RR, we only extract the intensity values and not the m/z nor the rt values. To compensate for this, we add the exact m/z and rt of the peptide from the corresponding MS1 scan as additional information saved in parallel with the other parts of the representation. This method is chosen to retain the locality of the peptide in MS1 space, while reducing data dimensionality for the CNN.

*MS2 level configuration.*

The MS2 spectrum representation, which can have Thermo peak picking applied to it (configurable in *config.json*), is extractable in two different ways, (1) the full spectrum with all m/z-intensity peaks (top n-peak filtering is required later on by ML applications to ensure conformity of input sizes) or (2) the spectrum split into bins of either fixed or variable sizes. Along with both of these methods, MS2AI selects the best matching MS2 spectra to any given MS1 peak (as multiple MS2 spectra can correspond to the same peptide). This task is automated in the MS2AI software, and is performed by matching precursor ion m/z of the available MS2 spectra to the peptide being matched with, then sorting the list to get the highest total intensity spectrum with the closest matched precursor-ion.

Once the best matching MS2 spectrum is found, the extraction method is applied. For the first method, the entire spectrum is retrieved from the mzML file and represented by a vector of peak pairings given by m/z and intensities (like the RR, the intensities are normalized between zero and one). The level of detail in this complete representation of the MS2 data might be prone to noise ultimately leading to a poor generalization of the ML models.

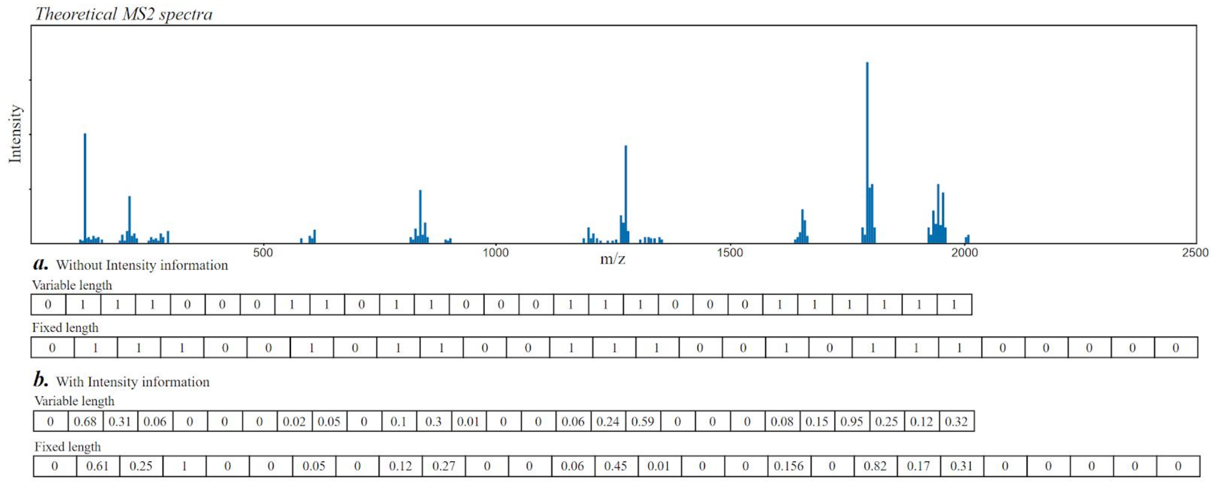


Figure 5. Illustration of the two methods used in MS2 spectra binning. This is illustrated using dummy values on a theoretical MS2 spectra.

The other method has two variations for binning the MS2 spectra. The parameters considered are: (a) number of bins, (b) variable or fixed lengths and (c) whether to retain intensity information. This method is illustrated in Figure 5 with 27 bins. Here the difference between variable and fixed lengths and whether to retain intensity information or not, is illustrated using an example MS2 spectrum with dummy values.

(a) *Number of bins:* Unlike the bin size of MS1, MS2AI does not directly divide MS2 data into m/z intervals, but rather takes a fixed number of bins, and superimposes the m/z interval that corresponds to the data. This parameter is user-specified in the *config.json* files, and defaults to 500.

(b) *Variable or fixed length:* As peptide masses often vary between ~200-2500 m/z, and similar ranges can be found in the MS2 spectra. For this reason we need to figure out how we should bin the MS2 spectra, whether to create a uniform method and bin all peptides into the same band of m/z, or to bin them between the highest and lowest m/z found in each given spectra. For example, if the peptide has an m/z of 2000, then binning these in 500 bins will result in each bin being 4 m/z wide, but if the peptide only has an m/z of 200, then each bin will be 0.4 m/z wide. This inconsistency in m/z representations could be an issue for ML applications, and for this reason we have chosen to default the binning method to a fixed m/z, this default length has been set to 2500. This will ensure a homogeneity of the m/z values in the MS2 data representation going into ML applications.

However, while we do gain homogeneity, we also create some potential issues. Peptides with lower m/z will have the important distinguishing factors located in the same bins, and thus they might be harder to distinguish , while having no information in the upper 450 bins.

(c) *Retain intensity values:* The intensities in an MS2 spectra can be further simplified by conversion into binary values indicating whether or not a bin contains at least one peak. With this method, a barcode-like representation of each peptide is created, potentially beneficial for ML applications for classification tasks with increased generalizability. On the other hand, dichotomizing the data discards a lot of potentially valuable information. Therefore, MS2AI provides an option using the average intensity of the peaks in each bin instead.

The data for our use case was done using MS2 binning, as this configuration gave the highest prediction. Table 2 shows a pros and cons list of binning vs non-binning.

| *MS2 Representations* | | | |
|---|---|---|---|
| **Full Information** | | **Binning** | |
| Pros | Cons | Pros | Cons |
| Retains all available information | Might not be generalizable. Retains too much knowledge | More generalizable | Discards a lot of intensity and peak data |
| Can utilize the differing MS2 scan lengths in ML applications | Requires specialized ML architecture to utilize differing length of MS2 scans | Variable bin sizes, allowing more user customizability | Hard to determine hyperparameters. Bins, length, etc. |
| | | Can retain some intensity information or create barcode-like data structure | |

Table 2. Pros and cons comparing the two methods of representing the MS2 spectra. The binning is combining pros and cons from variable and fixed size binning, as well as with retaining or discarding intensity binning.

#### c. Metadata Database

Regardless of the chosen extraction method, each peptide is accompanied by an entry in a MongoDB collection. This entry contains all information about the peptide taken from the MaxQuant output file specified in the configuration file together with the project specific information obtained by PRIDE or the local data repository. For instance, this project specific information could be the instrument used or the species the sample was taken from (find an overview of all keys in Table 1). This is important when it comes to filtering this database later on for specific ML applications.

When extracting peptides, the software will check for the unique ID of each peptide in the database, and skip it if this peptide already exists, even if it was created using different parameters for the creation of RR or PR. For this reason we have implemented a configurable database-name, allowing for multiple MongoDB databases to contain PR metadata, and thereby also the creation of multiple duplicates of the same PR (for separate projects, or different PR parameters). On Figure 2 this is expressed as the 'database-name' which is where the PRs are saved to allow for duplicates.
MS2AI also comes with a reset feature, that deletes all PRs along with their entries into the metadata collection. See *documentation.pdf* on the extractor for more information.

### 4. Data filtering

MS2AI comes with an inbuilt feature (see *documentation.pdf* on filter) to further filter the PR metadata and to ensure easy transformation into the most commonly used neural network package, TensorFlow[7].
The filter creates a new MongoDB collection in the given database, and only appends the following information to each entry. These information keys are: id, label, label name, project name (PXD), MS1 size, MS2 size, and name of filter class. This filtered version gives the individual subclasses (for example oxidation subclass within PTM class) numerical values according to the abundance found in the data, most abundant subclasses get lowest numerical values. This filtered version is used for the in-built neural network, and is specialized to work with a tailored data generator created in Keras[8].

An important parameter in the data filtering is the score filtering: Each peptide identified by MaxQuant has a corresponding score assigned ('Score' in MQ software). This score states how confident MaxQuant is in its assessment of the database search results; the lower the score, the lower the chance is that this peptide was classified correctly. The score filter calculates the distribution of scores in the PR metadata, and removes any entry with a score below the percentile stated by the user in the configuration file (calculated based on all PRs). Filtering based on score is applied to clean the noisy data, and remove potentially wrongly labeled peptides. Score thresholds are not generally comparable between different projects and should therefore be used with the necessary discretion.

When filtering the PR metadata, there are some parameters that need to be taken into consideration; (1) which class and subclass filter the data based on (Class: PTM, tissue. Subclass: oxidation, human), (2) number of classes to divide the data into, (3) subclass availability across all raw files, and (4) subclass imbalance handling.

(1) Depending on the class chosen for filtering, the data will be filtered into either a class (classification) or given a numerical value (regression). This is an automated process in the MS2AI software, where given a class, the software will either report back the numerical values (m/z, score, etc.) or a numerical value based on the abundance of subclasses in the class (for example filtering based on charge state, then charge state 2 will be given class 0, as it's the most abundant).

(2) Along with the class itself, an important parameter is also the amount of subclasses to include in the filtering. Currently MS2AI always sorts subclasses based on their abundance in the data, MS2AI will filter only the n-most abundant subclasses and discard data on everything else. This means that if users want to filter based on a low-abundance class they will either have to increase the subclass amount or use the binary subclass filtering also found in the software. Using the binary filtering feature, the user states the subclass of interest, and the MS2AI software labels everything within the subclass as '1', and everything not within the subclass as '0'. The two options allow for general exploratory research of different classes to take place, while also allowing research into specific details of different subclasses.

(3) MS identification softwares only performs database searches for things that the researchers are looking for, for example if a research paper is interested in phosphorylations, they will not perform database searches for a range of other PTMs, even though they might occur in the data, leading to mislabeled data. For this reason the user can also configure (in *config.json*) whether or not to filter data only from raw files where all of the subclasses occur, as this means that the search space for all given subclasses were taken into account during MaxQuant identification.

(4) Different subclasses have different distributions in the data, leading to different levels of frequency. This results in a class imbalance in the filtered data, meaning that there will be more data for certain subclasses than others. When it comes to class imbalance, MS2AI offers two separate options for solving this problem, either limiting the data to equal amounts of each subclass (chosen at random), or to report the class imbalance to Keras during network training, multiplying the reported accuracies with the class imbalance weights.
In addition, MS2AI offers the option to search through only the raw files where each of the classes are present

### 5. Use case neural network

Our PRs creates a concise version of the MS1 and MS2 data of a peptide, and to show the extent of the information retained in our PRs, MS2AI also comes with an example neural network with a tailored data generator in the neural network package, Tensorflow.

#### a. Data

For the use case we gathered ~6.5TB or ~69.000.000 PRs from 307 separate PRIDE projects. The 69m PRs were filtered into oxidized and not oxidized with a 98th percentile score filter. After filtering we had ~200.000 PRs for network training and testing. The filtered PRs were then further split into training (80% of data), validation (10% of data) and testing (10% of data) data; training and validation are taken from the same PRIDE projects, whereas testing is taken from entirely separate projects, in order to get as heterogeneous testing data as possible. The data was not filtered in any other regards, and the values used for the creation of PRs are the defaults found in *config.json*. These values are RR bins of 2 m/z and 0.2 rt, and PR neighbourhood intervals of 50 m/z and 5 rt.

The amount of data split into training, validating and testing are configurable parameters in the configuration file, and the process is entirely automated by MS2AI. Be advised, that testing projects are calculated based on when the amount of projects exceeds the amount requested in the configuration file. This means that if 5% of data is set as test data, then even if there is only one project, that entire project will be chosen as test data, and there will be no data labeled to train (as not doing so would cause there to be less than 5% test data). The split of testing data tries to be as accurate to the percentages requested, but it cannot be perfect as it removes entire projects and not just single PRs (the more data, the closer this separation will be).

#### b. Architecture

For MS1 we went with a convolutional layer of kernel size 5x5, and a max pooling layer of size 2x2. For the additional m/z and rt inputs from MS1 were handled by a single dense layer of size 64. The MS2 inputs were handled by a feed forward network with 3 layers of 256 neurons and dropout regularizers. These layers are subsequently concatenated and two dense layers of size 128 and 64 are added before a single neuron output layer. Testing to get to this network was in no way exhaustive, and can be changed by the user in the path/to/ms2ai/network/model.py script. The network is depicted in Figure 6.

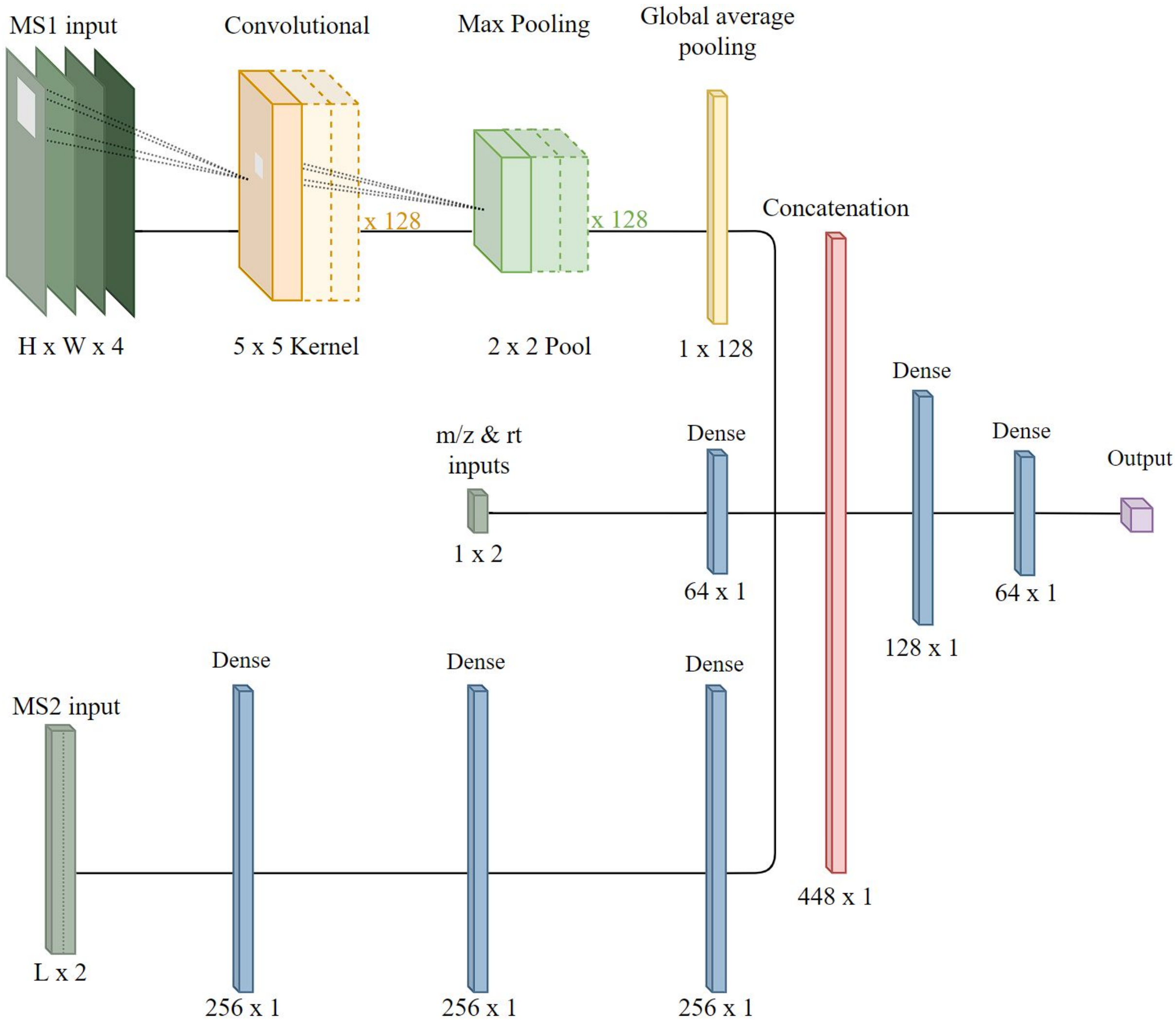


Figure 6. Neural network architecture used in the network training and testing for the paper. Also the architecture that comes with the software.

#### c. Metrics, cost functions and optimizers

During testing we attempted both classification and regressions problems, and for this we trained and tested with two main setups for metrics, cost functions and optimizers. For classifications we used binary or categorical cross entropy as our loss functions, with accuracy as our metric of success, and lastly we used adaptive momentum as the optimizer. For regression we used mean squared error as the loss function, r-squared as the metric for success, and lastly we used RMSprop as the optimizer.

Training was performed with a maximum of 50 epochs and with a 10 epoch early stopping (set a priori) in order to prevent overfitting. Furthermore, training was done on a batch size of 256 with a runtime per batch of ~2-3 seconds. Run time is predominantly defined by the size of data and the network; for more information on the runtime behavior refer to section 6.

All models use checkpoints to save the best model during training validated using validation accuracy, and the best is saved in the metadata folder on figure 2. The name scheme used for these files are the MS level followed by the MaxQuant software name for the predicted class, this could for example be ms1-Modification.h5 (for PTM classification), ms2-instrumentNames.h5 (for MS instrument classification) or both-Charge.h5 (for charge state classification). This naming scheme is important, as any pre-trained network has to be transferred into the metadata directory, and follow the correct naming scheme, in order to be processed by the MS2AI software.

##### d. Results

Distinguishing oxidized from non-oxidized peptides with an equal class balance, we obtained a training accuracy of 95%, a validation accuracy of 93% and a testing accuracy of 85%. This was achieved using a non-optimized neural network that mixes FNNs and CNNs for our different data inputs.

#### 6. Runtime evaluations

MS2AI allows a high amount of customizability when it comes to the data representation creation. However, changing the values from the defaults (default are in black text on Figure 7) can result in either much higher computational time needed for every step in both creation and ML process or oversimplification of the data. This section shows how such changes affect the computational time required for creating the RRs and the PRs, as well as the batch time for training the example neural network. The runtime calculations were done using a single raw file averaged over a few attempts for each file size.

| Peptide Creation time statistics | | | | | | | | | | NN Time statistics | |
|---|---|---|---|---|---|---|---|---|---|---|---|
| Full Scale Bins | | | | | Peptide Intervals | | | | | Batch (256) | |
| mz | rt | size | time | change | mz | int | size | time | change | time | change |
| 2 | 0.2 | 1 | 19.1 | | 50 | 5 | 250 | 15.8 | | 0.8 | |
| 2 | 0.2 | ↓1 | 19.1 | ↓100% | 100 | 10 | ↑1000 | 29.7 | ↑188% | 5 | ↑625% |
| 1 | 0.1 | ↑4 | 25.5 | ↑134% | 50 | 5 | ↑1000 | 43.8 | ↑277% | 5 | ↑625% |
| 1 | 0.1 | ↑4 | 25.5 | ↑134% | 100 | 10 | ↑4000 | 105.8 | ↑670% | 19.8 | ↑2475% |
| 5 | 0.5 | ↓0.16 | 14.9 | ↓78% | 10 | 2 | ↓3.2 | 6.1 | ↓39% | 0.03 | ↓4% |
| 0.5 | 0.05 | ↑16 | 41.1 | ↑215% | 100 | 10 | ↑16000 | 490 | ↑3101% | 90.3 | ↑11250% |

File amount: 3248

Table 3. Shows runtime calculations for 6 different calculations of full scale and PR creation and neural network training time. First (black) text is the baseline that the others are calculated based on.

It can be observed that changes affect the runtime dramatically, especially the training time of the neural networks. With a more complicated model than the one presented, the largest representations could potentially go much higher in computational time, and be completely unviable to train. This figure is important to keep in mind when wanting more complete data, as it comes with severe penalties in computational time.

#### 7. **Tutorial**

This section gives new users a short introduction to the general usage of MS2AI. This tutorial shows how to extract data from a pride project and apply the in-built neural network. For this reason the tutorial will skip the part of retrieving metadata from the PRIDE repository, as we do not use the metadata for the tutorial. (for a total documentation and usage tutorial see *documentation.pdf*)

The entire tutorial can be performed by running the following line in the terminal from the ms2ai directory

```
python api.py -T
Or to run the entire tutorial with individual tag entries
python api.py -e PXD019564 -f Modifications -b “Oxidation (M)” -n both -t
```

Each step will go through each cli option tag individually as it is used, and explain what it does.

##### a. Setup

This step isn't an option through the cli, it's a general setup of the configuration files and softwares. First the user needs to configure the path variable in *config.json* (see readme file for a detailed explanation for all configurable parameters), which needs to refer to an existing data folder. The default is ‘Data/’ from within the MS2AI directory. Next is the installation of the tools required by MS2AI. These are Python[9] 3.6+ (with pip), MongoDB[10], and either Conda[11] or Docker[12] for MacOS or Linux users, or only Docker for Windows users. The entire software is run via Python scripts. MongoDB has to be installed and a server instance needs to be available for MS2AI to work. Lastly, Docker or Conda is necessary for the required formatting tool, ThermoRawFileParser[1]. MacOS and Linux users can choose freely between the Conda and Docker, whereas Windows users have only the Docker choice. This tool can also be manually added via the code below in a terminal from the ms2ai directory. If these are not installed manually they will be installed during the first run of the software.

```
# Installation via Conda (a):
conda config --add channels defaults
conda config --add channels bioconda
conda config --add channels conda-forge
conda install -y -c bioconda thermorawfileparser

# Installation via Docker (b):
docker pull veitveit/trp
```

Along with the formatting software, the user also has to make sure that the formatting software specified in the configuration file corresponds correctly with the method installed.

##### b. Data extraction

The next step is to extract data from a given data source. For this tutorial we used a PRIDE project with only one small raw file (PXD019564) but in principle this tutorial also works with data from multiple projects.

For other options than running the extractor on single PRIDE projects, see *documentation.pdf.*

During this step, a directory for the project, and a sub directory for the raw files will be created in accordance to the folder structure described in Supplementary Figure 1. First the MaxQuant file is downloaded and cross-referenced with the raw file in the PRIDE database, then the corresponding raw file is downloaded, converted to mzML and discarded in order to save storage space. Next the information required to create the unique data representations are extracted from the converted file and stored for future usage. For the chosen PRIDE project 3248 PRs will be created. This step is performed by running the following line in the cmd or terminal from the ms2ai directory.

```
python api.py -e PXD019564
```

#### c. Filtering

The filtering process is used to prepare the data for a given machine learning task. In this tutorial, we seek to separate oxidized from non-oxidized peptides. Therefore, we configured MS2AI to use 'Oxidation (M)' as the active filter. When configuring filters, it is important to use the MaxQuant specific naming scheme such that MS2AI can identify the entries correctly. All available classes and their names can be found in the MaxQuant software documentation. Any peptide not classified by MaxQuant as 'Oxidation (M)' will be given the label 0, and peptides classified as 'Oxidation (M)' will be given the label 1.

Along with the oxidation filter, we also added data restriction to ensure that class balance. This data balance variable is stated in the *config.json* under the 'limit_data' key. No filter for the peptide confidence scores was added.

This filtering is performed by running the following line in the terminal from the ms2ai directory.

```
python api.py -f Modifications -b 'Oxidation (M)'
```

#### d. Machine learning application

The python script for this tutorial calls a pre-trained network located in the tutorial folder within the MS2AI gitlab directory. In order for the software to recognize this pre-trained network, this pre-trained model file *(tutorial.h5)* needs to be moved into the correct directory (metadata, see Figure 2) and renamed in accordance with the naming convention used for the project (here: *Best-Modifications-both.h5*). This naming convention includes the class used to filter (Modifications), and the levels of MS used for training (both MS1 and MS2).

Creating the directory and moving the model file is performed by running the following lines in a python interpreter, or by manually creating the needed paths and moving and renaming the tutorial.h5 file.

```
from storage.local import __create_file, __get_metapath
__create_file(__get_metapath())
shutil.copy('tutorial/tutorial.h5', __get_metapath() +
'Best-Modifications-both.h5')
```

Finally, we test the neural network with the in-built test function.

This is performed by the following line in the terminal from the ms2ai directory.

```
python ms2ai_api.py -n both -t
```

This script will test the data retrieved from the extraction of PXD019564 against the pre-trained network utilizing **both** MS1 and MS2 data (-n both), and reports back the test results. Due to the fact that only 3248 PRs were created during this tutorial, the network is only going to report 59.3% accuracy, as the test data amount is too low to converge to a solid average.

This concludes the tutorial. For more information read the *README.md* and *documentation.pdf*. The output of running the tutorial should be:

```
TUTORIAL: installing pre-required python packages
TUTORIAL: Retrieving data from pride accession PXD019564

Accessions: PXD019564
file: PXD019564/W8094TPRo_IP
W8094TPRo_IP: ✔
PXD019564 ✓. Total files: 1. Extracted files: 1.

TUTORIAL: Filtering data based on score and PTMs (oxidation)
```

```
Rough estimate of peptides to sort through: 3248
Rough estimate of runtime to filter: 0.01s or 0.0m
1: Oxidation (M) (114)
0: not Oxidation (M) (114)
Time elaborated: 0.101  | Per peptides (above reqs): 0.0  | Per peptides (all):
0.0

TUTORIAL: creating file system and running a neural network test on PTM data
train datapoints: 0
validation datapoints: 0
test datapoints: 228
1/1 [==============================] - 0s 1ms/step - loss: 1.3500 - accuracy:
0.5938
Accuracy on test data. Loss: 1.3499807119369507. Accuracy: 0.59375
```

```
📦/path/to/data
 ┣ 📂images
 ┃    ┗📂database-name
 ┃      ┗📂project-name
 ┃        ┗📂raw files
 ┃          ┗📜peptide-representation-ID.txt
 ┣ 📂metadata
 ┃    ┣📜best-model.h5
 ┃    ┣📜network-model.png
 ┃    ┗📜historic-plot.png
 ┣ 📂PXD projects
 ┃    ┗📂raw files
 ┃      ┣ 📜file.raw
 ┃      ┣ 📜file.mzML.gz
 ┃      ┣ 📜mzML.txt
 ┃      ┣ 📜MaxQuant-file.txt
 ┗      ┗ 📜XxY.txt
```
Supplementary Figure 1. Folder structure created by the MS2AI software at the path specified in the configuration file.
